# Aberrant functional brain network organization is associated with relapse during 1-year follow-up in alcohol-dependent patients

**DOI:** 10.1101/2023.06.01.543210

**Authors:** Justin Böhmer, Pablo Reinhardt, Maria Garbusow, Michael Marxen, Michael N. Smolka, Ulrich S. Zimmermann, Andreas Heinz, Danilo Bzdok, Eva Friedel, Johann D. Kruschwitz, Henrik Walter

## Abstract

Alcohol dependence (AD) is a debilitating disease associated with high relapse rates even after long periods of abstinence. Thus, elucidating neurobiological substrates of relapse risk is fundamental for the development of novel targeted interventions that could promote long-lasting abstinence. In the present study, we analyzed resting-state functional magnetic resonance imaging (rsfMRI) data from a sample of recently detoxified AD patients (*n* = 93) who were followed-up for 12 months after rsfMRI assessment. Specifically, we employed graph theoretic analyses to compare functional brain network topology and functional connectivity between future relapsers (REL, *n* = 59), future abstainers (ABS, *n* = 28) and age and gender matched controls (CON, *n* = 83). Our results suggest increased whole-brain network segregation, decreased global network integration and overall blunted connectivity strength in REL compared to CON. Conversely, we found evidence for a comparable network architecture in ABS relative to CON. At the nodal level, REL exhibited decreased integration and decoupling between multiple brain systems compared to CON, encompassing regions associated with higher-order executive functions, sensory and reward processing. Among AD patients, increased coupling between nodes implicated in reward valuation and salience attribution constitutes a particular risk factor for future relapse. Importantly, aberrant network organization in REL was consistently associated with shorter abstinence duration during follow-up, portending to a putative neural signature of relapse risk in AD. Future research should further evaluate the potential diagnostic value of the identified changes in network topology and functional connectivity for relapse prediction at the individual subject level.

## Introduction

Alcohol dependence (AD) is a debilitating disease with severe health consequences [1] and high relapse rates even after long periods of abstinence [2-4]. Though relapse rates vary considerably depending on the underlying definition of relapse and the employed follow-up interval, a recent meta-analysis based on twelve prospective studies with follow-up periods ranging from three to fifty years indicates that nearly every second patient relapsed between baseline and follow-up [5]. Thus, identification of neurobiological markers associated with relapse risk in AD is crucial to further the development of targeted interventions that facilitate long-lasting abstinence and reduce health burdens associated with AD.

Resting-state functional magnetic resonance imaging (rsfMRI) has revealed aberrant resting-state functional connectivity (rsFC) in AD patients in general [6-9] and in relapsing AD patients in particular [10-14]. For instance, Camchong et al. (2022) reported decreased rsFC within four large-scale functional brain networks associated with executive control, incentive salience and negative emotionality in subsequent relapsers compared to future abstainers. Importantly, decreased rsFC within each functional network predicted subsequent time to relapse during a 4-months follow-up period [10]. However, these previous studies either investigated alterations of predefined large-scale functional brain networks [10] or employed seed-based approaches to probe deviations in rsFC associated with relapse in AD [11-14], limiting findings to a narrow set of seed regions and network definitions.

Graph theory, in turn, provides a data-driven approach to characterize the topological organization of functional brain networks [15]. At the level of the whole brain, network segregation and network integration delineate two fundamental concepts in functional connectomics that provide insights into specialized information processing within subgroups of interconnected brain regions (segregation) and the efficiency of information propagation across the brain (integration) [16]. At the level of individual brain regions, centrality metrics indicate the differential importance of single nodes for the entire network [16]. Previous research investigating brain network topology in AD using graph theory revealed increased global clustering [17], decreased global efficiency [18], reduced average functional connectivity and fragmentation of functional modules [19] in alcohol-dependent patients compared to healthy controls, indicating more segregated and less integrated global network topology in AD with overall decreased connectivity strength. Probing brain-behavior relationships, Sjoerds et al. (2017) found that reduced global clustering, reduced global efficiency and reduced network degree were associated with longer and more severe alcohol dependence [20]. However, research on the topological features of rsFC networks associated with relapse risk in AD is scarce. One study investigated the impact of transcranial direct current stimulation (tDCS) on brain network topology in alcohol use disorder (AUD) and its association to relapse [21]. Specifically, AUD patients underwent tDCS over the bilateral dorsolateral prefrontal cortex for five consecutive days and participated in rsfMRI sessions before and after the intervention. Afterwards, they were followed-up for 90 days to assess alcohol relapse. Global efficiency significantly increased from baseline to post-tDCS, while global clustering decreased. Importantly, increased global efficiency predicted reduced likelihood of early relapse during follow-up while decreased global clustering was associated with reduced impulsivity [21], highlighting aberrant network segregation and network integration as a potential marker for relapse risk in alcohol-dependent individuals.

In the present study, we therefore investigated the resting-state functional brain network architecture in a sample of recently detoxified alcohol-dependent patients who were followed up for 12 months using graph theory. Specifically, we compared measures of whole-brain network organization (network strength, network segregation, network integration), local network organization (nodal centrality) and inter-regional functional connectivity (Network-Based Statistics, NBS [22]) between future relapsers (REL), future abstainers (ABS) and controls (CON) to determine topological network characteristics associated with relapse risk in AD. At the whole-brain level, we hypothesized that particularly REL would display increased network segregation, decreased network integration and overall decreased connectivity strength as compared to CON. At the level of individual nodes, we expected decreased centrality and functional connectivity of brain regions and networks previously implicated in AD relapsers, including regions associated with higher-order executive control functions, reward processing, sensory processing and negative emotionality [10-13]. Conversely, we expected comparable network organizations in ABS and CON, as previous work suggests [23]. Finally, we assumed that putative disruptions in network topology and functional connectivity in REL would relate to abstinence duration during follow-up, suggesting a potential neural substrate of relapse risk in AD.

## Methods

### Participants

All subjects were recruited as part of a large, bicentric study conducted in Berlin and Dresden investigating predictors for the development and maintenance of alcoholism (ClinicalTrials.gov identifier: NCT01679145). Data of this study has been reported in previous work addressing different research questions [e.g., 24-26]. In total, 222 subjects enrolled in this study. After quality control, 177 subjects with valid behavioral and imaging data were included in the present sample. Ninety-three patients aged 18 to 65 suffering from alcohol dependence for at least three years according to ICD-10 and DSM-IV-TR criteria (*M* = 11.62 years according to DSM-IV-TR, *SD* = 10.05) were assessed shortly after their detoxification treatment (*M* = 23.23 days, *SD* = 12.63). Additionally, 84 age and gender matched controls with social drinking behavior were included in the study as a control group. Patients were excluded if they reported a current or past substance use disorder except for alcohol and nicotine, other major psychiatric disorders, neurological disorders or psychoactive medication. For relapse assessment, AD patients were continuously followed-up for 12 months (48 weeks) after the MRI measurement using the Timeline-Follow-Back (TLFB [27]), a reliable method to retrospectively assess alcohol use [28]. Particularly, subjects were a) contacted via telephone 6, 10, 18 and 36 weeks after the last occasion of alcohol use, and b) assessed in person 4, 8, 12, 24 and 48 weeks after the last drinking occasion. Follow-up assessments were completed by 87 AD patients (93.5%). Relapse was defined as one drinking occasion during the follow-up interval with at least 60 g or 48 g of alcohol intake for male and female subjects, respectively, or when the phosphatidylethanol (PEth) levels determined from dried blood spots of venous blood at either 12- or 24-weeks follow-up exceeded 112 ng/ml [29]. Consequently, our sample consisted of 59 relapsers (REL) and 28 abstainers (ABS), see **Tab. 1**. The study was carried out in line with the Declaration of Helsinki and was approved by the medical ethics committees of the Charité – Universitätsmedizin Berlin and Technische Universität Dresden. All subjects provided informed written consent prior to study participation.

**Table 1.**
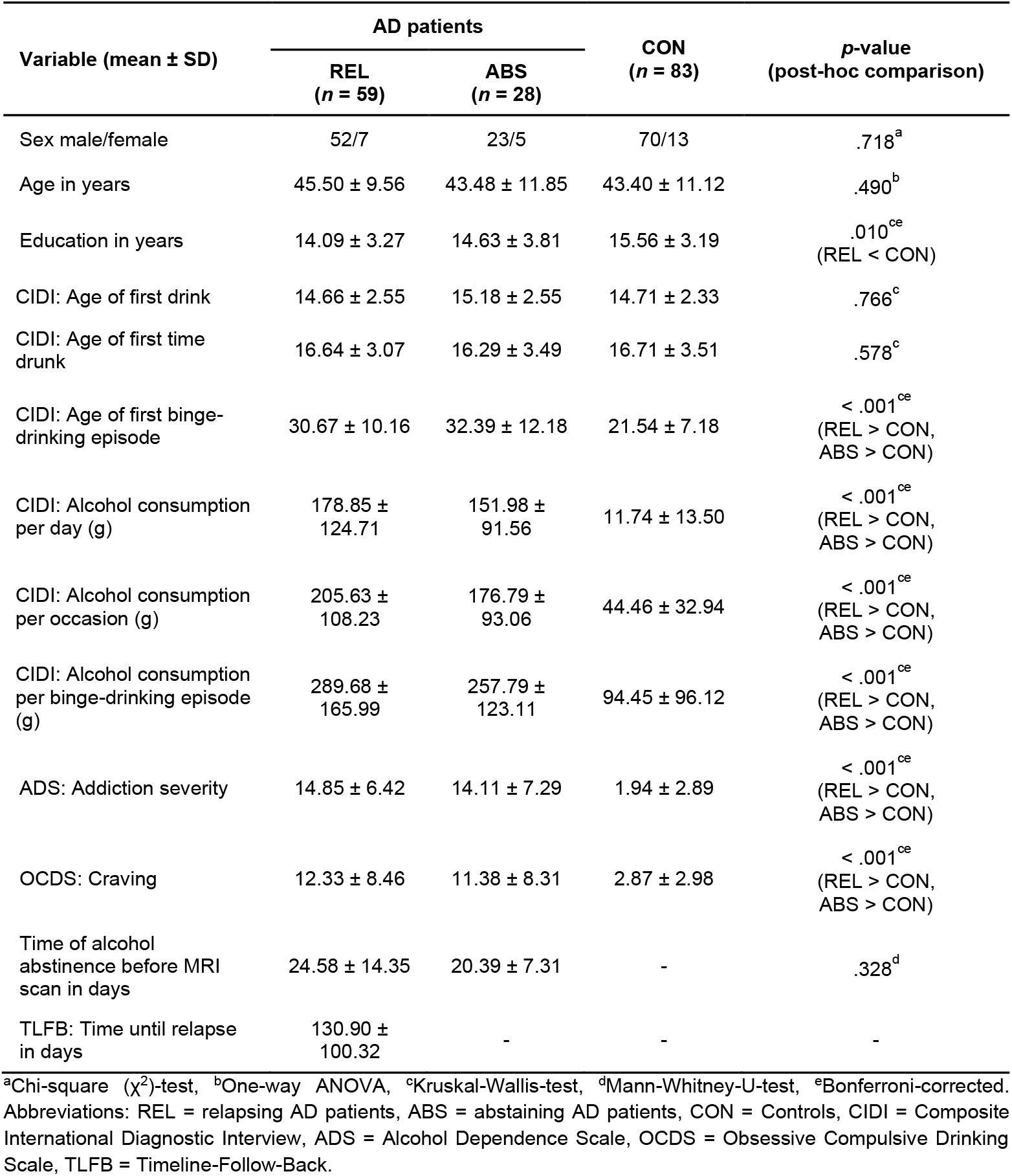
Demographics and clinical characteristics.

### Clinical and psychometric assessment

For diagnosis and assessment of individual drinking behavior, we used the computerized version of the Composite International Diagnostic Interview (CIDI [30,31]). AD severity was assessed using the Alcohol Dependence Scale (ADS [32]). Moreover, we implemented the Obsessive Compulsive Drinking Scale (OCDS [33]) to measure alcohol craving and we recorded the number of days of abstinence before MRI assessment. For REL, we also assessed the number of days between the MRI measurement and the first relapse using the TLFB.

### Image acquisition

All images were acquired on two 3T Siemens Magnetom Tim Trio MRI scanners in Berlin and Dresden with a 12-channel phased-array head coil (Siemens Healthineers, Erlangen, Germany), respectively. For each participant, 148 functional images were acquired during resting-state using a gradient-echo echo-planar imaging (GE-EPI) sequence (TR = 2410 ms, TE = 25 ms, 80 ° flip angle, 42 axial (ax > cor) slices with 1 mm slice gap, 2 mm slice thickness, FOV = 192×192 mm^2^, 3×3 mm^2^ in-plane resolution, bandwidth = 2112 Hz/Px). During data acquisition, participants were asked to lie as still as possible while keeping their eyes closed without thinking of anything specific or falling asleep. Additionally, high-resolution T1-weighted 3D structural images (MPRAGE, TR = 1900 ms, TE = 2.52 ms, 9° flip angle, isotropic voxel size of 1×1×1 mm^3^, bandwidth = 170 Hz/Px) and a B0-field map (TR = 488 ms, TE = 5.32 ms, 60° flip angle, isotropic voxel size of 3×3×2 mm^3^, bandwidth = 260 Hz/Px) were acquired.

### Data preprocessing

Data preprocessing was performed using FSL v5.0 [34], including motion correction, acquisition delay correction, non-brain tissue removal, spatial low-pass filtering with an isotropic gaussian kernel of 6 mm FWHM, denoising using ICA-based artifact removal (ICA-AROMA [35]), high-pass filtering of the time series with a cut-off frequency of 0.008 Hz, co-registration of resting-state volumes to the T1 image using boundary-based registration with a field map based correction of spatial distortions due to local field inhomogeneity, non-linear normalization of the T1 image to the 2 mm MNI standard space template (Montreal Neurological Institute, Quebec, Canada) using Advanced Normalization Tools (ANTs [36]), spatial transformation of resting-state data to MNI standard space applying the registration matrices and warp images from the two previous registration steps, and resampling of resting-state data into 3 mm isotropic voxels. Participants with excessive head movements after head motion correction (i.e., mean framewise displacement > 0.5 mm) were excluded.

### Network construction

#### Node definition

Nodes were obtained by dividing the brain into anatomically distinct regions of interest (ROI) based on the Brainnetome Atlas [37], resulting in a network of 246 nodes (210 cortical and 36 subcortical nodes). Each node’s time series was calculated as the mean of the time series from all voxels belonging to the respective node.

#### Edge definition

We estimated functional connectivity between nodes using L2-regularized partial correlation (regularization parameter *rho* = 0.01), calculated via the inverse of the covariance matrix. This resulted in a 246×246 rsFC matrix for each participant. Partial correlation determines the correlation between two node’s time series while conditioning on the time series from all other nodes in the network, thus reflecting variance uniquely shared between two nodes, above and beyond ups and downs in functional connectivity that would be shared with other network nodes (in the linear modeling regime) [38]. In this way, partial correlation produces more accurate estimates of direct connectivity as compared to naïve, raw node-node correlation estimates [39,40], making it a pertinent method to investigate biologically informed alterations of network topology in disease populations [41].

### Network topology analysis

To reduce the possibility of bias due to a specific connection threshold, we applied a set of sparsity thresholds to the partial correlation matrices using a proportional threshold range of 0.10 < *k* < 0.50 with intervals of 0.01, where *k* specifies the ratio of strongest edges that are maintained in the network relative to the maximum number of possible edges in the network. Thresholding can introduce network fragmentation, i.e., splitting up the network into disconnected components where at least one node is not connected to any other node in the network, thereby confounding the computation of network topological parameters. In our study, we assessed network fragmentation for all subjects and thresholds using the function CheckFrag.m implemented in GraphVar 2.0 [42,43].

The topological properties of the undirected weighted network were calculated for each threshold, including global measures on network segregation (global clustering coefficient *C*_*global*_, global transitivity *T*_*global*_), network integration (global efficiency *E*_*global*_, global characteristic path length *L*_*global*_) and global connectivity strength *W*_*global*_. At the nodal level, we evaluated the topological centrality of each node as an indicator for its influence on the network. To this end, we calculated four different measures of nodal centrality, including nodal degree centrality *D*_*nodal*_, nodal strength centrality *W*_*nodal*_, nodal eigenvector centrality *EC*_*nodal*_ and nodal betweenness centrality *BC*_*nodal*_. A detailed description of the global and nodal graph measures analyzed in this study can be found in **Tab. S1**. Prior to the computation of graph topological metrics (except for *W*_*global*_), we scaled subject-level rsFC matrices by the maximum absolute weight to obviate confounds in network topology introduced through potential differences in weight distributions. To improve interpretability, subject-level graph metrics were additionally normalized by dividing the respective measure of the original network by the average of the same measure derived from 100 random networks with the same number of nodes and similar degree, weight and strength distributions (see function null_model_und_sign.m from Brain Connectivity Toolbox Version 2019-03-03; [16]; available at https://sites.google.com/site/bctnet/all-functions). For each normalized global and nodal graph measure, we computed the area under the curve (AUC) as an integrated score independent of a single threshold selection. Network analyses were performed using GraphVar 2.0 [42,43] running under the MATLAB^®^ environment.

### Statistical analysis

#### Group differences in global and local network topology

Group comparisons of global and nodal graph metrics between AD subgroups (REL, ABS) and CON were carried out using nonparametric permutation tests controlling for age, gender, site and education years. First, we computed the one-way ANOVA *F*-value testing for differences in the AUC of a given graph metric between groups. Next, we randomly shuffled the group labels between participants and re-calculated the *F*-value. We repeated this procedure 10,000 times to create an empirical null distribution of the effect underlying group differences in AUC. Finally, we obtained a *p*-value by estimating the proportion of permutations that were equal to or exceeded the empirical *F*-value. Results were considered significant if actual effects exceeded those from the empirical null distribution according to the *p* < .05 threshold. Global and nodal graph measures exhibiting significant group differences were further subjected to post-hoc analysis. Specifically, we computed the empirical differences in AUC between groups using two-sample t-tests, then permuted the group labels 10,000 times, re-calculated the t-test and estimated the proportion of effects in the permutation-based null distribution that exceeded the empirical group effect. Results were considered significant if *p* < .05. A false discovery rate (FDR) correction accounting for multiple testing was implemented for global and nodal metrics separately. For global metrics, we accounted for the number of global parameters and post-hoc group comparisons (*n* = 15) while for nodal metrics, we corrected for the number of parameters (*n* = 246) for each graph metric. To assess evidence for the null hypothesis, we also calculated the Bayes factor for each group contrast based on t-tests with a JZS prior (*r* = √2/2) using the BayesFactor package in R [44] and scaled the corresponding values using a log10 transform.

#### Group differences in functional connectivity

We applied the Network-Based Statistic (NBS) approach [22] to the fully connected partial correlation matrices to identify differences in rsFC between groups. Briefly, significantly altered subnetworks, defined in terms of connected graph components, were determined by incorporating an edge-by-edge comparison of rsFC based on a one-way ANOVA with *F* > 4.8 (*p* < .01). Within the identified subnetworks, post-hoc two-sample t-tests were conducted to detect significantly altered components between specific groups. In our study, we used an initial link threshold of *t*_*initial*_ = 3.1 to identify suprathreshold links. Since NBS highly depends on the initial cluster-defining threshold, we additionally tested a range of different link thresholds (*t*_*initial*_ = 2.9-3.3). To estimate the significance of the identified components, we generated an empirical null distribution of maximal component sizes by randomly permuting the group assignments 10,000 times and calculating the proportion of permutations in which the maximal component size exceeded the size of the observed component. Components (i.e., subnetworks) were considered significant if *p* < .05, FWE-corrected.

#### Brain-behavior relationships

To investigate associations between putative alterations in network topological parameters and time to first relapse to alcohol use in AD patients, we used partial correlation controlling for age, gender, site and education years. In case of non-normality of clinical and graph topological variables as determined by Shapiro-Wilk tests (*p* < .05), we computed Spearman’s partial rank correlation. FDR correction determined over the number of significantly altered network metrics was applied to account for multiple testing.

## Results

### Sample characteristics

One subject was excluded from further analyses due to severe network fragmentation. As a result, 59 relapsing AD patients (REL), 28 abstaining AD patients (ABS) and 83 control subjects (CON) were included in the final analyses (**Tab. 1**). No significant differences in age and sex distributions were observed in any of the group comparisons. REL reported fewer years of education compared to CON. As expected, both AD patient subgroups reported significantly more alcohol consumption (per day, per drinking occasion and per binge-drinking episode), higher addiction severity (ADS) and higher alcohol craving (OCDS) compared to CON. Interestingly, CON experienced their first binge-drinking episode at an earlier age compared to both AD patient subgroups. No significant differences could be observed between REL and ABS on any demographic or clinical outcome (**Tab. 1**).

### Global topology of functional brain networks

REL exhibited decreased *W*_*global*_ (*p*_*FDR*_ = .042), increased *C*_*global*_ (*p*_*FDR*_ = .004), increased *T*_*global*_ (*p*_*FDR*_ = .004), decreased *E*_*global*_ (*p*_*FDR*_ = .039) and increased *L*_*global*_ (*p*_*FDR*_ = .039) compared to CON, suggesting a more segregated and less integrated network topology in REL with overall decreased network strength (**Fig. 1A**). No significant differences were found between ABS and REL as well as between ABS and CON on any graph measure after FDR correction. Bayes Factor indices based on t-tests corroborated results from frequentist analyses. Specifically, we found moderate to strong evidence for differences between REL and CON across all global graph measures, while no such evidence was found for the comparison of REL vs. ABS (**Tab. S2**). Moreover, in line with our hypotheses, comparing ABS vs. CON consistently yielded anecdotal or moderate evidence for the null hypothesis across all global graph measures (**Tab. S2**).

**Figure 1.**
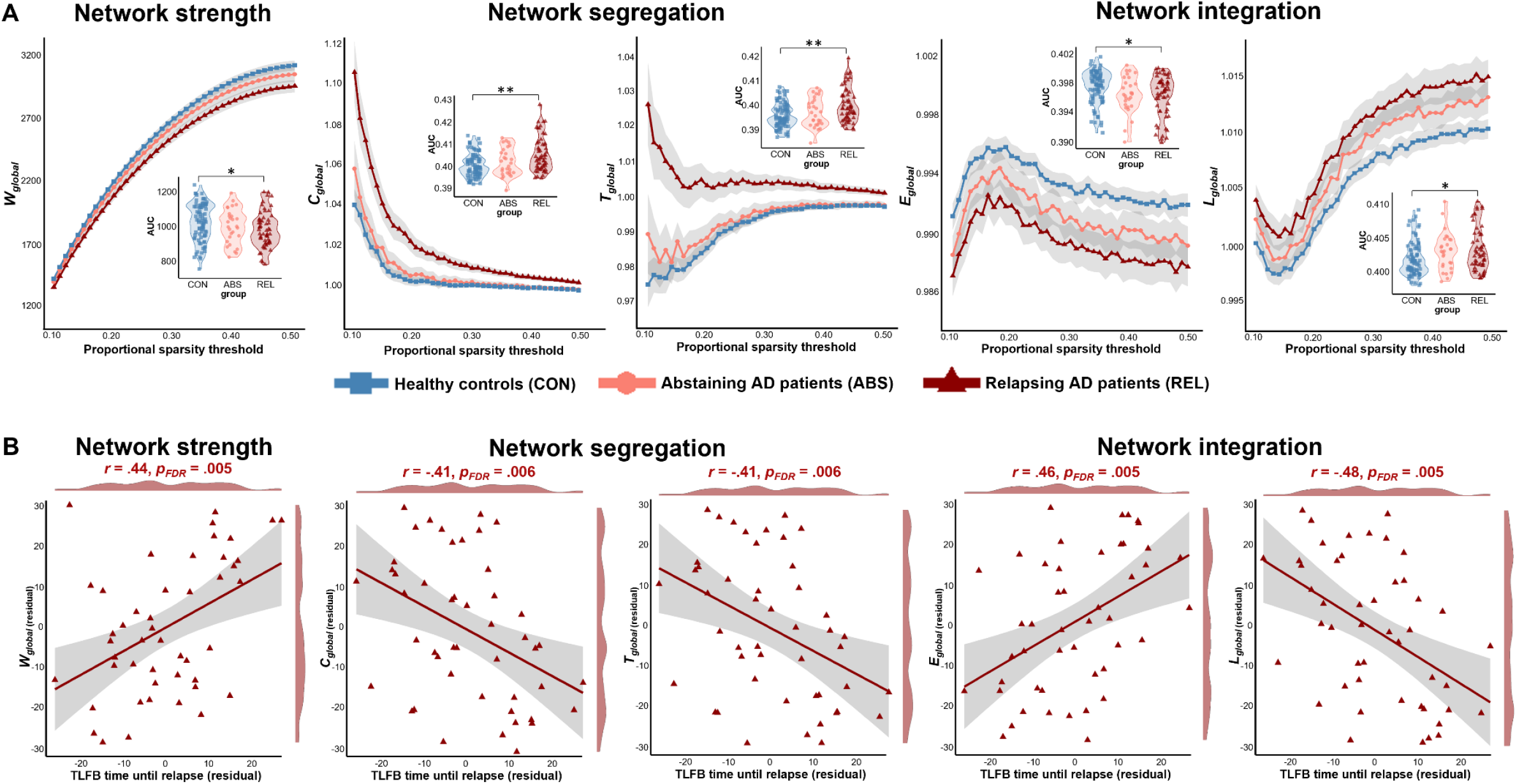
A) Global network organization in relapsing AD patients (REL, dark red), abstaining AD patients (ABS, light red) and controls (CON, blue), indicating decreased network strength and more segregated and less integrated network topology in REL compared to CON. B) Partial correlations (controlling for age, gender, site and education years) between altered graph measures in REL and the time until the first relapse (measured using Timeline-Follow-Back, TLFB) show that earlier relapse is associated with decreased network strength, increased network segregation and decreased network integration. \**p* < 0.05, \*\**p* < 0.01 after FDR correction.

### Nodal centrality

At the nodal level, we identified those brain regions that displayed significant group differences based on frequentist analysis (*p* < .05, 10,000 permutations, FDR-corrected) as well as strong evidence for group differences based on Bayes Factor indices (*BF*_*log(10)*_ > 1) for any nodal graph measure. In this regard, REL exhibited decreased nodal strength centrality (*W*_*nodal*_) in 12 cortical and ten subcortical brain regions compared to CON. The strongest evidence for reduced nodal strength centrality was observed in the left putamen (**Tab. 2**). Moreover, REL displayed decreased nodal eigenvector centrality (*EC*_*nodal*_) in five subcortical brain regions compared to CON, with the strongest evidence for group differences in the left ventromedial putamen (**Tab. 2**). We found no differences between REL and CON in degree centrality and betweenness centrality. Comparing REL vs. ABS and ABS vs. CON yielded no differences in any nodal centrality measure, respectively.

**Table 2.** Brain regions exhibiting altered nodal centrality in relapsing AD patients (REL) compared to controls (CON).

| Brain region | | Nodal strength<br>($W_{nodal}$) | | | Nodal eigenvector centrality<br>($EC_{nodal}$) | | |
| --- | --- | --- | --- | --- | --- | --- | --- |
| Brainnetome region | Anatomical structure | AUC difference | $p$-value<sup>a</sup> | Bayes Factor<sup>b</sup> | AUC difference | $p$-value<sup>a</sup> | Bayes Factor<sup>b</sup> |
| **Frontal regions** |  |  |  |  |  |  |  |
| SFG_L_7_6 | BA 9 medial | -0.29 | .049* | 1.14<sup>†</sup> | -0.0008 | .351 | 0.34 |
| IFG_L_6_5 | BA 44 opercular | -0.33 | .044* | 1.89<sup>††</sup> | -0.0010 | .307 | 0.51 |
| PCL_R_2_1 | BA 1/2/3 lower limb region | -0.26 | .047* | 1.17<sup>†</sup> | -0.0004 | .633 | -0.29 |
| **Temporal regions** |  |  |  |  |  |  |  |
| FuG_R_3_1 | BA 20 rostroventral | -0.24 | .044* | 1.25<sup>†</sup> | -0.0009 | .378 | -0.02 |
| FuG_R_3_2 | BA 37 medioventral | -0.26 | .047* | 1.22<sup>†</sup> | -0.0007 | .462 | -0.02 |
| PhG_L_6_1 | BA 35/36 rostral | -0.32 | .049* | 1.05<sup>†</sup> | -0.0007 | .604 | -0.25 |
| PhG_R_6_2 | BA 35/36 caudal | -0.34 | .047* | 1.29<sup>†</sup> | -0.0008 | .492 | -0.18 |
| PhG_R_6_3 | lateral posterior parahippocampal gyrus | -0.34 | .044* | 2.07<sup>††</sup> | -0.0009 | .164 | 1.67<sup>††</sup> |
| PhG_L_6_6 | medial posterior parahippocampal gyrus | -0.33 | .047* | 1.22<sup>†</sup> | -0.0004 | .680 | -0.43 |
| **Limbic regions** |  |  |  |  |  |  |  |
| CG_L_7_4 | BA 23 ventral | -0.35 | .047* | 1.47<sup>†</sup> | -0.0009 | .295 | 0.74 |

| **Occipital regions** |  |  |  |  |  |  |  |
| --- | --- | --- | --- | --- | --- | --- | --- |
| MVOcC_L_5_3 | caudal cuneus gyrus | -0.33 | .047* | 1.46<sup>†</sup> | -0.0010 | .461 | 0.04 |
| MVOcC_R_5_4 | rostral lingual gyrus | -0.28 | .047* | 1.22<sup>†</sup> | -0.0007 | .555 | -0.12 |
| **Subcortical regions** |  |  |  |  |  |  |  |
| Amyg_L_2_2 | lateral amygdala | -0.35 | .047* | 1.14<sup>†</sup> | -0.0010 | .351 | 0.10 |
| Hipp_L_2_1 | rostral hippocampus | -0.28 | .049* | 1.29<sup>†</sup> | -0.0008 | .512 | 0.08 |
| BG_L_6_2 | globus pallidus | -0.35 | .044* | 1.69<sup>††</sup> | -0.0015 | .041* | 2.14<sup>†††</sup> |
| BG_R_6_2 | globus pallidus | -0.32 | .044* | 1.73<sup>††</sup> | -0.0012 | .105 | 1.62<sup>††</sup> |
| BG_L_6_4 | ventromedial putamen | -0.40 | .020* | 2.42<sup>†††</sup> | -0.0017 | .041* | 4.01<sup>†††</sup> |
| BG_R_6_4 | ventromedial putamen | -0.36 | .047* | 1.72<sup>††</sup> | -0.0012 | .041* | 1.99<sup>††</sup> |
| BG_L_6_6 | dorsolateral putamen | -0.36 | .020* | 2.48<sup>†††</sup> | -0.0014 | .041* | 2.28<sup>†††</sup> |
| BG_R_6_6 | dorsolateral putamen | -0.29 | .049* | 1.50<sup>††</sup> | -0.0009 | .316 | 1.22<sup>†</sup> |
| Tha_L_8_2 | pre-motor thalamus | -0.39 | .020* | 2.13<sup>†††</sup> | -0.0015 | .041* | 2.41<sup>†††</sup> |
| Tha_R_8_8 | lateral prefrontal thalamus | -0.23 | .047* | 1.04<sup>†</sup> | -0.0007 | .378 | -0.09 |

<sup>a</sup>*p*-value based on 10,000 permutations (FDR corrected); <sup>b</sup>Bayes Factor (*BF*) based on t-tests with a JZS prior ( $r = \sqrt{2/2}$ ), rescaled using log10 transform. \**p* < 0.05. <sup>†</sup> $BF_{\log(10)} 1 - 1.48$ (strong evidence for H1), <sup>††</sup> $BF_{\log(10)} 1.48 - 2$ (very strong evidence for H1), <sup>†††</sup> $BF_{\log(10)} > 2$ (extreme evidence for H1) based on a heuristic classification scheme by Schönbrodt and Wagenmakers (2018) [45]. Abbreviations: BA = Brodmann area, SFG = superior frontal gyrus, IFG = inferior frontal gyrus, PCL = paracentral lobule, FuG = fusiform gyrus, PhG = parahippocampal gyrus, CG = cingulate gyrus, MVOcC = medioventral occipital cortex, Amyg = amygdala, Hipp = hippocampus, BG = basal ganglia, Tha = thalamus, L = left, R = right.

### Brain-behavior relationships

Aberrant global brain network organization in REL significantly correlated with the duration of abstinence during follow-up (**Fig. 1B**). Specifically, earlier relapse was associated with reduced global network strength (*W*_*global*_: *r* = .44, *p*_*FDR*_ = .005) as well as more segregated (*C*_*global*_: *r* = -.41, *p*_*FDR*_ = .006; *T*_*global*_: *r* = -.41, *p*_*FDR*_ = .006) and less integrated brain networks (*E*_*global*_: *r* = .46, *p*_*FDR*_ = .005; *L*_*global*_: *r* = -.48, *p*_*FDR*_ = .005). At the nodal level, earlier relapse in REL was associated with decreased nodal strength centrality in the left dorsomedial prefrontal cortex (SFG_L_7_6), right paracentral lobule (PCL_R_2_1), right inferior temporal gyrus (FuG_R_3_2), right parahippocampal gyrus (PhG_R_6_2, PhG_R_6_3), left posterior cingulate gyrus (CG_L_7_4), bilateral globus pallidus (BG_L_6_2, BG_R_6_2) and left pre-motor thalamus (Tha_L_8_2) (**Fig. 2**).

**Figure 2.**
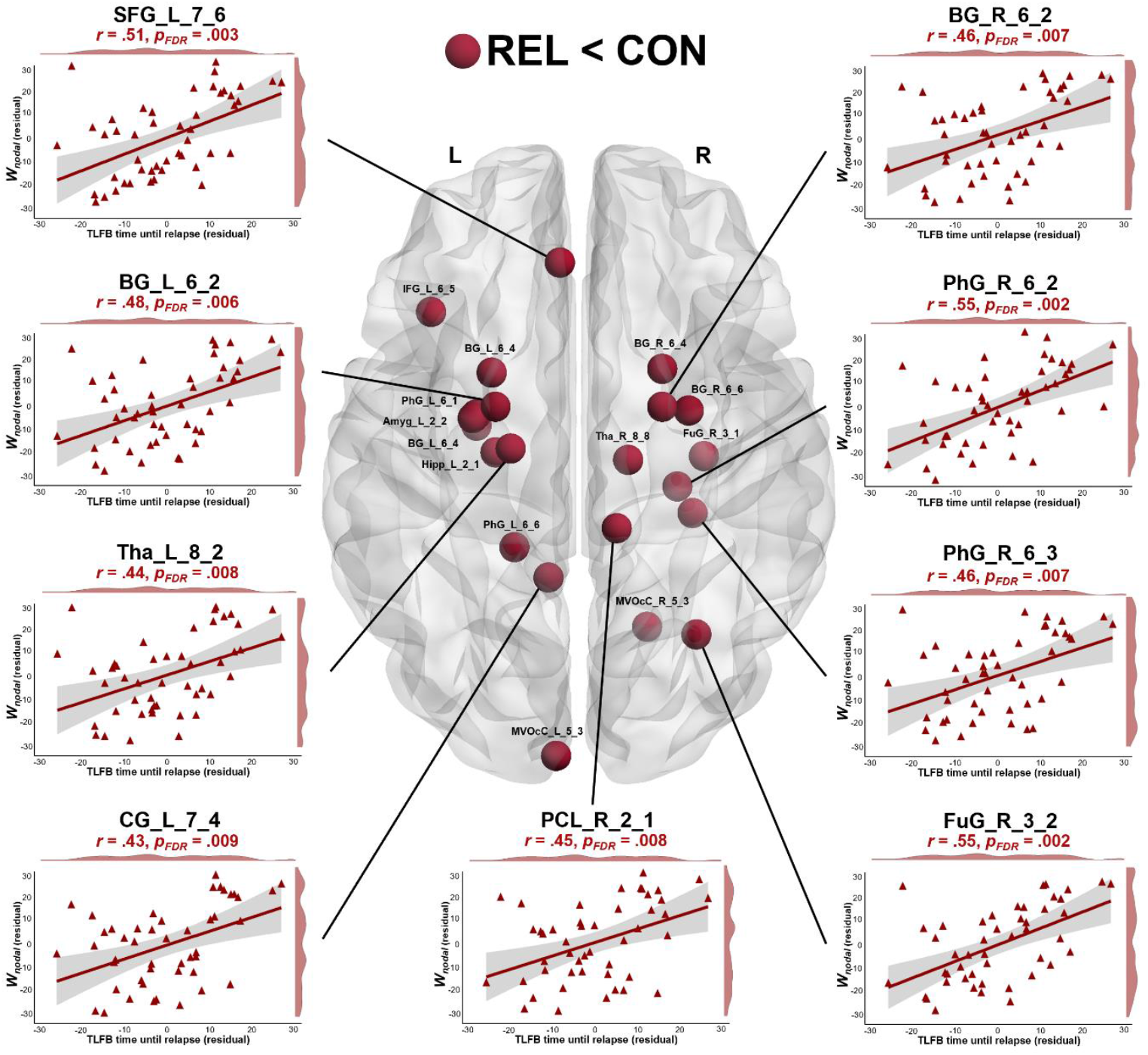
Brain regions exhibiting reduced nodal centrality (*W*_*nodal*_, *EC*_*nodal*_) in relapsing AD patients (REL) compared to controls (CON). Decreased *W*_*nodal*_ in left dorsomedial prefrontal cortex (SFG_L_7_6), right paracentral lobule (PCL_R_2_1), right inferior temporal gyrus (FuG_R_3_2), right parahippocampal gyrus (PhG_R_6_2, PhG_R_6_3), left posterior cingulate gyrus (CG_L_7_4), bilateral globus pallidus (BG_L_6_2, BG_R_6_2) and left pre-motor thalamus (Tha_L_8_2) in REL correlated with the time until the first relapse occurred (based on Timeline-Follow-Back, TLFB) as revealed by partial correlation analysis controlling for age, gender, site and education years.

### Network-based statistics

NBS analysis revealed a predominantly left-hemispheric subnetwork of significantly decreased rsFC in REL compared to CON (*p*_*FWE*_ = .015), comprising eight nodes and seven edges. Specifically, REL exhibited decreased rsFC mainly between the precuneus (PCun_L_4_1) and the dorsomedial prefrontal cortex (SFG_L_7_6), orbitofrontal cortex (OrG_L_6_3), lateral occipital cortex (LOcC_L_4_1) and the lateral prefrontal thalamus (Tha_L_8_8), as well as between lateral occipital cortex (LOcC_L_4_1) and parahippocampal and superior parietal cortex (PhG_L_6_1, SPL_R_5_4), and between dorsolateral precentral gyrus (PrG_L_6_2) and lateral prefrontal thalamus (Tha_L_8_8) (**Fig. 3A**). The largest difference in connectivity was observed between the precuneus and lateral occipital cortex (**Tab. 3**). Alternating the cluster-defining link threshold yielded similar subnetworks of significantly decreased connectivity in REL compared to CON (**Fig. S1**).

**Figure 3.**
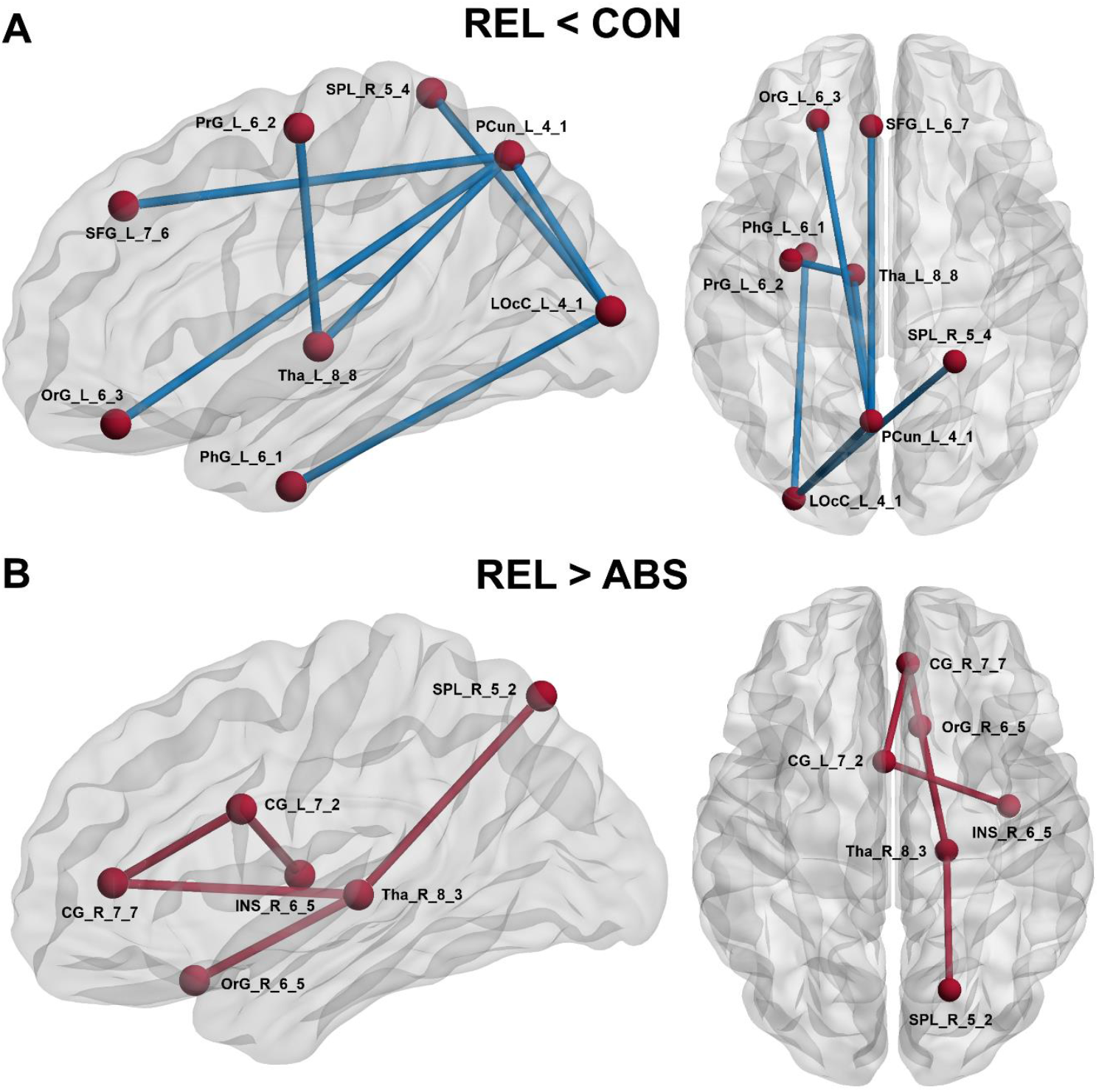
Network-based statistics (NBS) analysis revealed A) a subnetwork of decreased functional connectivity (blue) in REL compared to CON and B) a subnetwork of increased functional connectivity (red) in REL compared to ABS (10,000 permutations, initial link threshold *t*_*initial*_ = 3.1). REL = relapsing AD patients, ABS = abstaining AD patients, CON = controls.

**Table 3.** Edges exhibiting significantly altered functional connectivity in relapsing AD patients compared to controls (REL < CON) and in relapsing AD patients compared to abstaining AD patients (REL > ABS) as identified by NBS.

| Group contrast | Subnetwork edge | Difference in mean rsFC | t-value | p-value |
| --- | --- | --- | --- | --- |
| REL < CON | SFG_L_7_6 – PCun_L_4_1 | -0.0181 | 3.70 | <.001 |
|  | OrG_L_6_3 – PCun_L_4_1 | -0.0154 | 3.19 | <.001 |
|  | PhG_L_6_1 – LOcC_L_4_1 | -0.0165 | 3.38 | <.001 |
|  | SPL_R_5_4 – LOcC_L_4_1 | -0.0185 | 3.27 | <.001 |
|  | PCun_L_4_1 – LOcC_L_4_1 | -0.0191 | 4.04 | <.001 |
|  | PrG_L_6_2 – Tha_L_8_8 | -0.0135 | 3.52 | <.001 |
|  | PCun_L_4_1 – Tha_L_8_8 | -0.0137 | 3.18 | <.001 |
| REL > ABS | INS_R_6_5 – CG_L_7_2 | 0.0259 | 3.14 | .001 |
|  | CG_L_7_2 – CG_R_7_7 | 0.0249 | 3.97 | <.001 |
|  | OrG_R_6_5 – Tha_R_8_3 | 0.0262 | 3.31 | <.001 |
|  | SPL_R_5_2 – Tha_R_8_3 | 0.0248 | 3.26 | <.001 |
|  | CG_R_7_7 – Tha_R_8_3 | 0.0260 | 3.19 | <.001 |

Abbreviations: SFG = superior frontal gyrus, Pcun = precuneus, OrG = orbital gyrus, PhG = parahippocampal gyrus, LOcC = lateral occipital cortex, SPL = superior parietal lobule, PrG = precentral gyrus, Tha = thalamus, INS = insula, CG = cingulate gyrus, L = left, R = right.

Moreover, we found a predominantly right-hemispheric subnetwork of significantly increased connectivity in REL compared to ABS (*p*_*FWE*_ = .036), comprising nodes in the dorsal anterior cingulate cortex (dACC, CG_L_7_2), subgenual anterior cingulate cortex (sgACC, CG_R_7_7) anterior insula (INS_R_6_5), orbitofrontal cortex (OFC, OrG_R_6_5), superior parietal lobule (SPL, SPL_R_5_2) and sensory thalamus (Tha_R_8_3) (**Fig. 3B**), with the largest difference in rsFC between the thalamus and orbitofrontal cortex (**Tab. 3**). The subnetwork replicated across different initial link thresholds but reached statistical significance only at *t*_*initial*_ = 3.1 (**Fig. S2**).

## Discussion

Despite extensive research on the chronically relapsing nature of AD, little is known about the topological features of functional brain networks related to relapse. Here, we investigated resting-state functional network topology and functional connectivity in future relapsers (REL) and future abstainers (ABS) assessed during a 12-months follow-up interval to identify neurobiological substrates associated with relapse risk in AD. Compared to controls (CON), REL displayed disturbances in global and local brain network topology, implicating multiple regions from different brain circuits. Conversely, ABS exhibited a comparable network organization relative to CON. Importantly, topological network characteristics in REL were associated with an earlier relapse after detoxification, portending to a possible neural signature of relapse risk in AD.

Examining the whole-brain global network topology, we observed decreased global functional connectivity strength in REL compared to CON, reminiscent of previous work [10,11]. Moreover, graph theoretic indices of functional network segregation revealed increased global clustering and global transitivity in REL compared to CON, suggesting more segregated brain networks in future relapsers relative to controls. It has been proposed that network segregation delineates the extent of specialized information processing within clusters of functionally related brain regions [16]. Elevated segregation, however, results in isolated information processing within subnetworks, thus perturbing the dispersion of information across the network [46]. Consistent with this view, we found impairments in global functional integration as indicated by increased global characteristic path length and decreased global efficiency in REL, suggesting diminished capacity to efficiently propagate information from distributed brain regions across the network. Together, our findings reveal a tendency for more segregated information processing at the expense of long-range cortical interactions between distributed brain regions, possibly reflecting impairments in cognitive flexibility typical for AD [47,48] that manifest in mental and behavioral rigidity, such as craving and compulsive alcohol use, and contribute to increased relapse risk [49]. Accordingly, aberrant network topological metrics were associated with shorter duration of abstinence during follow-up. Likewise, Holla et al. (2020) demonstrated that changes in global clustering and global efficiency decreased impulsivity and reduced likelihood of early relapse in AUD patients following tDCS treatment [21].

Similar alterations in global network organization have been shown for AD patients independent of relapse status [17-19], although results were not consistent across studies [20]. The present study extends previous work by highlighting deviations in network topology particularly in future relapsers. Conversely, we did not observe any significant differences in network topology between ABS and CON. In fact, Bayesian analyses consistently indicated more evidence for a comparable network organization in ABS and CON than evidence for different network topologies, giving rise to the conclusion that functional segregation and integration are largely preserved in those AD patients that prospectively refrain from further alcohol use. Of note, we did not observe any significant differences between REL and ABS with respect to alcohol use patterns, addiction severity or craving, debilitating the ascription of the observed effects to more severely affected individuals. Instead, the identified changes seem to be specific to the relapsing subtype, portending to a possible neural substrate of relapse risk in AD.

At the level of individual nodes, REL exhibited decreased centrality of several widely distributed regions implicated in multiple brain systems, many of which correlated with shorter abstinence duration after detoxification. In line with our predictions, we found decreased nodal strength centrality in prefrontal brain regions, including the left dorsomedial prefrontal cortex (dmPFC, BA 9) and ventrolateral prefrontal cortex (vlPFC, BA 44), two critical neural substrates involved in a variety of higher-order executive functions, including inhibitory control, self-regulation and goal-directed behavioral control [50,51]. Blunted vlPFC activation [52] as well as decreased dmPFC connectivity across different cognitive tasks [13,53] have been linked to relapse in previous work, substantiating the link between decreased regulatory control and relapse in AD. Moreover, NBS analysis revealed decreased rsFC in REL compared to CON between the dmPFC and the precuneus, a core region of the default mode network (DMN [54]). The DMN plays an important role for internally oriented cognitions, such as self-awareness, episodic memory and reflecting on past and future events [55]. Decoupling between the DMN (precuneus) and executive control network (dmPFC) has previously been observed in addiction [56], reflecting difficulties to disengage from negative emotions and ruminations about alcohol use during withdrawal [57], a known risk factor for relapse [58].

Furthermore, local network analyses revealed decreased centrality in two other core regions of the DMN, namely the posterior cingulate cortex (PCC, BA 23 [59]) and parahippocampal gyrus (PhG [60]). Given the DMNs role in self-reflection and memory processes, decreased centrality in PCC and PhG might indicate deficits in relapsers to reflect on past events of alcohol use and their consequences, thus compromising the capacity to adapt future decision-making behavior accordingly. Consistent with this view, decreased DMN node centrality was associated with earlier relapse after detoxification, extending previous findings of a positive relationship between efficiency in DMN nodes and length of abstinence [6]. In addition, consistent with our hypotheses, we observed decreased centrality of sensorimotor (BA 1,2,3) and visual network nodes (BA 20, BA 37, lingual gyrus, cuneus) in the whole-brain network. Hypoconnectivity in visual and sensorimotor domains have been related to alcohol-drinking [61-63] and AD [9,11] in previous studies. Decoupling between these two systems in relation to relapse was also demonstrated in the present study using the NBS approach. Recent work revealed an association between sensorimotor and visual network connectivity with trait impulsivity [64]. Thus, our findings indicate deficient functioning of brain systems in REL that support the integration of multimodal perceptual information, resulting in compromised sensory awareness that is relevant for decision-making processes, including the decision to inhibit alcohol use urges [11].

Importantly, relapse to alcohol consumption was associated with decreased centrality in several subcortical brain regions known to be critically involved at different stages of the addiction cycle, including the amygdala, hippocampus, basal ganglia (globus pallidus, putamen) and thalamus [65-67]. Consistent with our findings, Bordier et al. (2021) demonstrated a fragmentation of the basal functional module comprising the amygdala, pallidum, putamen, hippocampus, thalamus, and caudate into smaller subdivision in AD patients, which still persisted after two weeks of withdrawal treatment [19], suggesting a potential risk factor for perpetuated alcohol seeking. The strongest evidence for decreased centrality in REL was observed in the left putamen, a part of the dorsal striatum, which has been implicated in the transition from voluntary towards compulsive and habitual drug seeking [68]. The putamen functionally connects to motor cortices and prefrontal control regions, including the dmPFC and vlPFC [69,70]. Decreased putamen connectivity in REL might reflect the dysregulation of prefrontal control over the dorsal striatum that may propel motor systems towards habitual alcohol seeking. In this regard, impaired functional connectivity between dorsal striatum and prefrontal regions has been reported in cannabis dependent individuals at rest [71] and in AD patients when making habitual decisions, the latter being associated with shorter abstinence duration [72].

Interestingly, NBS analysis revealed that REL exhibited increased rsFC relative to ABS in a predominantly right-hemispheric subnetwork encompassing nodes in the ACC (dACC, sgACC), anterior insula, OFC, SPL and thalamus. The dACC and anterior insula delineate core regions of the salience network, guiding attentional resources towards salient events based on interoceptive states [73,74], while the sgACC and OFC constitute integral parts of the reward network which is implicated in the appraisal of subjective value [75,76]. Previous studies demonstrated greater OFC and ACC activation in response to alcohol-related cues in AD patients which was associated with alcohol craving and relapse [77-79]. Similarly, anterior insula function has recently been proposed as a marker for relapse propensity in AD [80] due to its role for craving and drug seeking [81]. The thalamus acts as a hub region in the identified subnetwork with functional connections to the reward system nodes (sgACC and OFC) and to the SPL, a brain region also implicated in reward processing [82] and goal-directed allocation of attentional resources [83] that gets activated by alcohol cues in dependent drinkers [84]. Together, our findings indicate that hyperconnectivity between regions implicated in reward processing and salience attribution contributes to relapse risk among AD patients, possibly reflecting elevated preoccupation and anticipation of reward during the brain’s resting state that may propel individuals towards alcohol-seeking behavior and thus facilitate relapse risk. These individuals might particularly benefit from behavioral interventions targeted at the modification of cognitive biases contributing to habitual behaviors [85].

Some limitations in this study should be considered. Firstly, we obtained a relatively small sample size in the group of AD abstainers. However, high relapse rates constitute a core characteristic of AD [5], thus lower sample sizes of abstainers relative to relapsers lie in the nature of the disorder. Nonetheless, future studies should employ larger samples to increase statistical power. Secondly, subjects were not matched based on socioeconomic status even though previous work highlighted the impact of socioeconomic factors for AD [e.g., 86]. However, in our study, we included information on the educational background of individuals as covariates when performing group analyses to control for confounding effects attributable to socioeconomic influences. Thirdly, the diagnostic value of the identified changes in functional network organization for the prediction of relapse at the level of the individual subject remains unknown [87,88]. Although previous studies demonstrated substantial predictive utility of functional network measures for individual diagnosis and classification of alcohol dependent individuals [62], future research should validate the identified pattern of functional connectome disorganization for individual relapse prediction.

In sum, the present study reveals distinct functional connectome disturbances in prospectively relapsing AD patients. Particularly, on a whole-brain level, relapsers process information rather in a segregated fashion within functionally interconnected regions than distributing information efficiently across the brain. On a local level, relapsers displayed dysfunctional integration and decoupling between multiple brain systems, particularly involving cortical and subcortical brain regions associated with higher-order executive functions, sensory and reward processing. Among AD patients, hyperconnectivity between brain regions implicated in reward processing and salience attribution seems to constitute a particular risk factor for relapse. Together, our findings provide novel insights into the neurobiological substrates of relapse risk in AD. Future research should further investigate the predictive utility of the identified pattern of functional connectome disorganization for relapse at the single-subject level.

## Acknowledgements

This work was supported by the German Research Foundation (Deutsche Forschungsgemeinschaft, DFG project numbers 186318919 [FOR 16717 to AH, EF, MNS, HW]; 178833530 [SFB 940 to MNS, MM, HW]; 402170461 [TRR 265 to AH, MNS, MM, HW]; 454245598 [GRK 2773 to MNS]) as well as the German Federal Ministry of Education and Research (Bundesministerium für Bildung und Forschung, BMBF project ID 01ZX1909C [SysMedSUDs to HW, AH]). The funders had no role in the study design, data collection, analysis, decision to publish or preparation of the manuscript.

## Author contributions

AH, MNS, USZ and HW were involved in the planning and conceptualization of the study. JB performed statistical analyses. JB drafted the manuscript. HW, JDK, EF, PR, MG, MM, MNS, USZ, AH, DB provided critical revision of the manuscript for important intellectual content. All authors critically reviewed the content of the manuscript and approved the final version for publication.

## Data availability statement

The dataset of this study is available upon request. The data are not publicly available due to privacy or ethical restrictions.

