## Supplemental Materials for "Aberrant functional brain network organization is associated with relapse during 1-year follow-up in alcohol-dependent patients"

Table S1. Description of global and nodal network measures investigated in this study, based on Rubinov and Sporns (2010).

| **Network level** | **Network measure** | **Calculation** | **Explanation** |
| --- | --- | --- | --- |
| Global network measures | Strength (*W_global_*) | $Wglobal= \sum_{i,j\in N} wij$  *N* = network, *i,j* = 1…*N*-1 nodes, $wij$ = connection weight between node *i* and *j* | The sum of all edge weights in the network. |
|  | Clustering coefficient (*C_global_*) | $Cglobal= \frac{1}{n}\sum_{i\in N} \frac{2tiw}{ki (ki-1)}$  *N* = network, *n* = number of nodes in *N*, *i* = node, $ki$ = degree of node *i*, $tiw$= geometric mean of triangles around node *i* | The ratio of triangles (=neighbours of a node that are also connected to each other) around a node relative to the maximum possible number of triangles around that node, making it a node-wise normalized measure on network segregation. The global clustering coefficient denotes the average clustering coefficient across all nodes in the network. |
|  | Transitivity (*T_global_*) | $Tglobal= \frac{\sum_{i\in N} 2tiw}{\sum_{i\in N} ki(ki-1)}$  *N* = network, *i* = node, $ki$ = degree of node *i*, $tiw$= geometric mean of triangles around node *i* | The ratio of triangles in the network relative to the total number of connected triplets in the network. In contrast to the clustering coefficient, transitivity represents a network-wise normalized measure on network segregation. |
|  | Characteristic path length (*L_global_*) | $Lglobal= \frac{1}{n} \sum_{i\in N} \frac{\sum_{j\in N,j\neq i} dijw}{n-1}$  *N* = network, *n* = number of nodes in *N*, *i* = node, *j* = 1…*N*-1 nodes, $dijw$= shortest weighted path length between nodes *i* and *j* | The average shortest path length between all possible node pairs in the network. A short characteristic path length allows more rapid information transfer across the network. Characteristic path length thus serves as an indicator for the network’s integrational ability. |
|  | Efficiency (*E_global_*) | $Eglobal= \frac{1}{n} \sum_{i\in N} \frac{\sum_{j\in N,j\neq i} {(dijw)}^{-1}}{n-1}$  *N* = network, *n* = number of nodes in *N*, *i* = node, *j* = 1…*N*-1 nodes, $dijw$= shortest weighted path length between nodes *i* and *j* | The average inverse of the shortest path length. In contrast to the characteristic path length, global efficiency is less influenced by the presence of disconnected nodes in the network. |
| Nodal network measures | Degree centrality (*D_nodal_*) | $Dnodal(i)= \sum_{j\in N} aij$  *N* = network, *i* = node, *j* = 1…*N*-1 nodes, $aij$ = connection status between node *i* and *j* with *a*$ij$ = 1 when a link exists and *a*$ij$ = 0 when no link exists | The total number of edges connected to a node. |
|  | Strength centrality (*W_nodal_*) | $Wnodal(i)= \sum_{j\in N} wij$  *N* = network, *i* = node, *j* = 1…*N*-1 nodes, $wij$ = connection weight between node *i* and *j* | The sum of weights from all edges connected to a node. |
|  | Eigenvector centrality (*EC_nodal_*) | $ECnodal\left( i \right)=$ $\frac{1}{\lambda1} \sum_{j\in N} Aij\chi j$  *N* = network, *i* = node, *j* = 1…*N*-1 nodes, $A$ = adjacency matrix, $\lambda1$ = largest eigenvalue of $A$, $\chi j$= eigenvector of $A$ for node *j* | The summed centrality of a node’s neighbours. Nodes have high eigenvector centrality when they are connected to nodes which themselves are highly connected. |
|  | Betweenness centrality (*BC_nodal_*) | $BCnodal\left( i \right)= \frac{1}{\left( n-1 \right)\left( n-2 \right)} \sum_{\begin{aligned} h,j\in N \\ h\neq j,h\neq i,j\neq i \end{aligned}} \frac{\rho hj(i)}{\rho hj}$  $\text{N}\text{ = network, }\text{n = }\text{number of nodes in }\text{N}\text{,}\text{ i}\text{ = node,}\text{ h,j}\text{ = 1…}\text{N}\text{-1 nodes,} \rho hj$ = number of shortest paths between *h* and *j*, $\rho hj(i)$ = number of shortest paths between *h* and *j* that pass through *i* | The ratio of shortest paths between all node pairs in the network that pass through a node. |

Table S2. Comparison of global network organization in relapsing AD patients (REL), abstaining AD patients (ABS) and controls (CON).

| **Post-hoc  group contrast** | **Graph measure^a^** | | **AUC difference** | ***p-*value^b^** | **Bayes Factor^c^** |
| --- | --- | --- | --- | --- | --- |
| REL vs. ABS | network strength | *W_global_* | -26.3183 | .484 | -0.36 |
|  | network segregation | *C_global_* | 0.0042 | .140 | 0.01 |
|  |  | *T_global_* | 0.0037 | .140 | 0.01 |
|  | network integration | *E_global_* | -0.0007 | .490 | -0.49 |
|  |  | *L_global_* | 0.0008 | .490 | -0.49 |
| REL vs. CON | network strength | *W_global_* | -46.4946 | **.042** | 0.60 |
|  | network segregation | *C_global_* | 0.0051 | **.004** | **1.52** |
|  |  | *T_global_* | 0.0045 | **.004** | **1.52** |
|  | network integration | *E_global_* | -0.0015 | **.039** | 0.74 |
|  |  | *L_global_* | 0.0016 | **.039** | 0.58 |
| ABS vs. CON | network strength | *W_global_* | -20.1764 | .490 | -0.50 |
|  | network segregation | *C_global_* | 0.0009 | .531 | -0.56 |
|  |  | *T_global_* | 0.0008 | .531 | -0.57 |
|  | network integration | *E_global_* | -0.0008 | .306 | -0.25 |
|  |  | *L_global_* | 0.0008 | .332 | -0.31 |

^a^Global clustering coefficient *C_global_*, global transitivity *T_global_*, global efficiency *E_global_* and global characteristic path length *L_global_* were normalized by the corresponding measure from 100 random networks with equal number of nodes and similar weight and strength distributions; ^b^*p-*value based on 10,000 permutations (FDR correction applied); ^c^Bayes Factor based on t-tests with a JZS prior (*r* = √2/2), rescaled using log10 transform. Results indicating *p* < .05 and BF_log10_ > 1 (strong evidence for H1) are highlighted in bold.

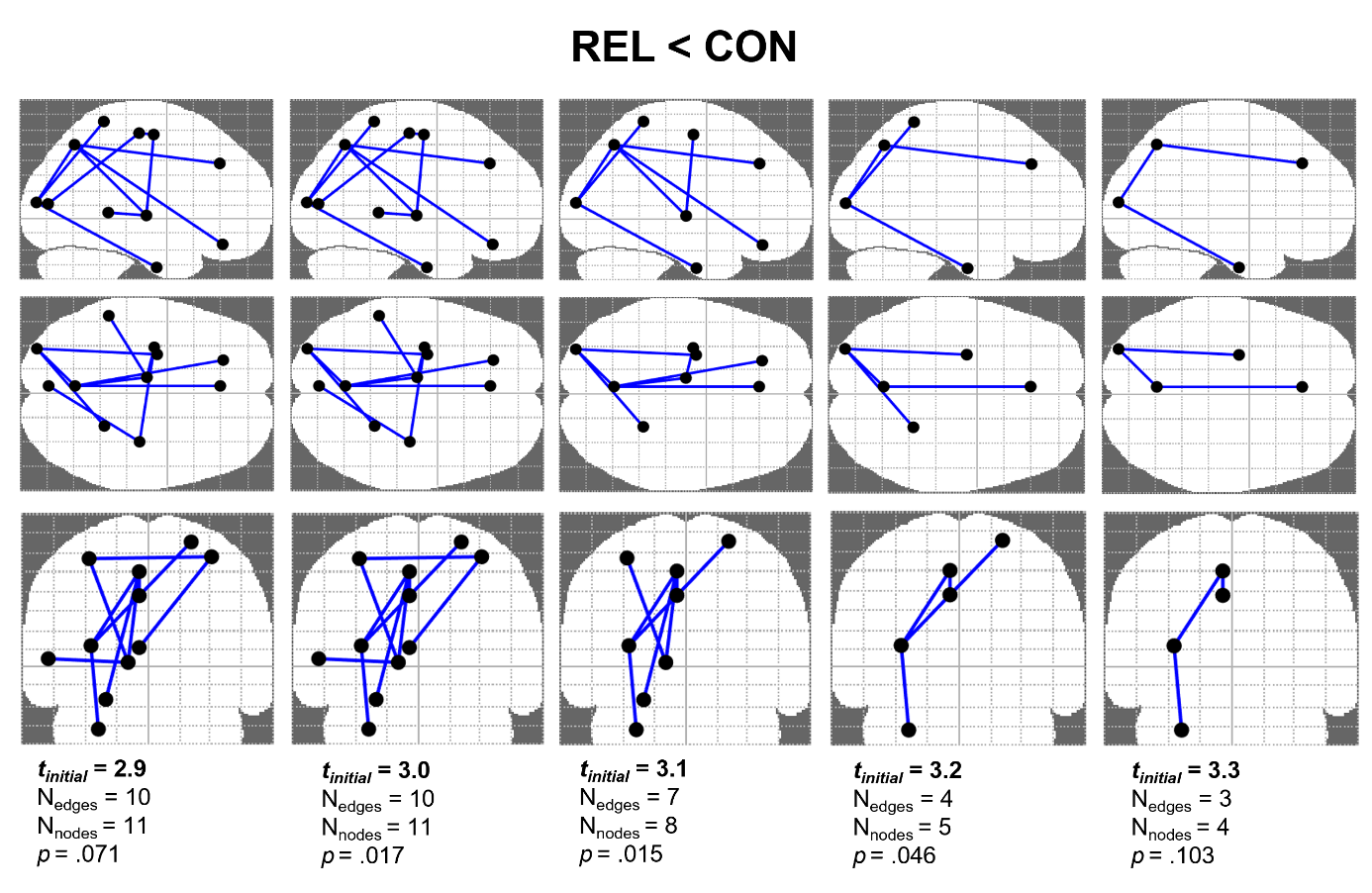

Figure S1. Network-based statistics (NBS) analysis comparing functional connectivity between relapsing AD patients (REL) and controls (CON) with alternating initial cluster-defining thresholds (*t_initial_* = 2.9-3.3). Results consistently reveal a subnetwork of decreased functional connectivity in REL compared to CON across different statistical thresholds. In the manuscript, we report the median link threshold (*t_initial_* = 3.1, corresponds to *p* = .001). *p*-values are based on 10,000 permutations.

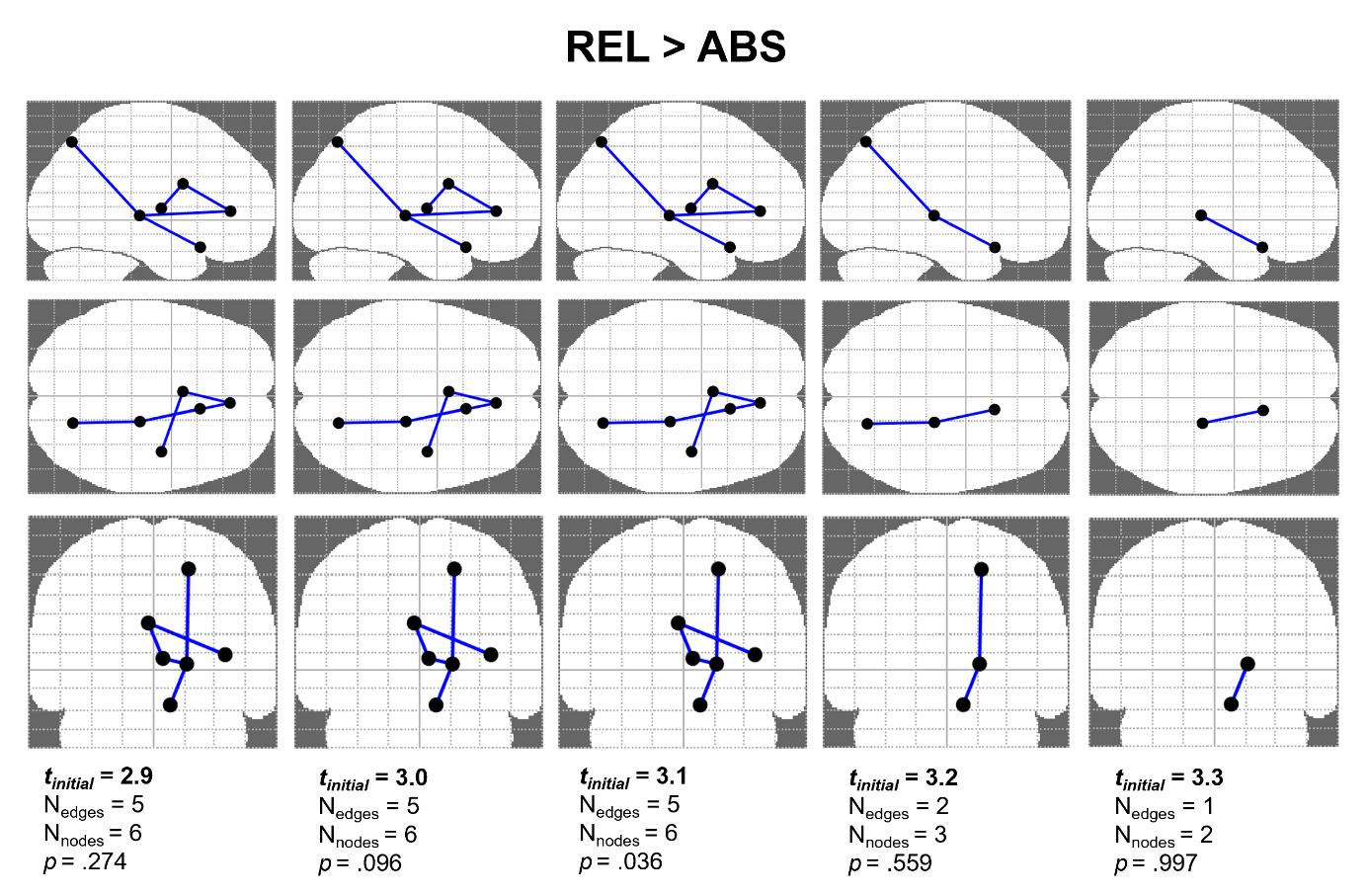

Figure S2. Network-based statistics (NBS) analysis comparing functional connectivity between relapsing (REL) and abstaining AD patients (ABS) with alternating initial cluster-defining thresholds (*t_initial_* = 2.9-3.3). Results reveal a subnetwork of significantly increased functional connectivity in REL compared to ABS at the median link threshold *t_initial_* = 3.1 (corresponds to *p* = .001). *p*-values are based on 10,000 permutations.
